# Metagenomic signature of natural strongyle infection in susceptible and resistant horses

**DOI:** 10.1101/233015

**Authors:** Allison Clark, Guillaume Sallé, Valentine Ballan, Fabrice Reigner, Annabelle Meynadier, Jacques Cortet, Christine Koch, Mickaёl Riou, Alexandra Blanchard, Núria Mach

**Affiliations:** Health Science Department, Open University of Catalonia, Barcelona, Spain; UMR 1282, INRA, Infectiologie et Santé Publique and Université François-Rabelais, Nouzilly, France; UMR 1313, INRA, AgroParisTech, Université Paris-Saclay, Jouy-en-Josas, France; UEPAO 1297, INRA, Unité Expérimentale de Physiologie Animale de l’Orfrasière, Nouzilly; UMR 1388, INRA, GenPhySE, Toulouse, France; UE-1277, INRA, Plate-Forme d’Infectiologie Expérimentale, Nouzilly, France; Pancosma SA, CH-1218 Geneva, Switzerland

**Author notes:** These authors contributed equally to this work (Allison Clark, Guillaume Sallé). Corresponding Author: Núria Mach Mail.

**Keywords:** cyathostomin, fungi, gut microbiome, immunity, horse

## Abstract

Gastrointestinal strongyles are a major threat to horses' health and welfare. Given that strongyles inhabit the same niche as the gut microbiota, they may interact with each other. These beneficial or detrimental interactions are unknown in horses and could partly explain contrasted susceptibility to infection between individuals. To address these questions, an experimental pasture trial with 20 worm-free female Welsh ponies (10 susceptible (S) and 10 resistant (R) to parasite infection) was implemented for five months. Fecal egg counts (FEC), hematological and biochemical data, body weight and gut microbiota composition were studied in each individual after 0, 24, 43, 92 and 132 grazing days.

The predicted R ponies exhibited lower FEC after 92 and 132 grazing days, and showed higher levels of circulating monocytes and eosinophils, while S ponies developed lymphocytosis by the end of the trial. Although the overall microbiota diversity remained similar between the two groups, R and S ponies exhibited sustained differential abundances in *Clostridium XIVa, Ruminococcus, Acetivibrio* and unclassified *Lachnospiracea* at day 0. These bacteria may hence contribute to the intrinsic pony resistance towards strongyle infection. Moreover, *Paludibacter, Campylobacter, Bacillus, Pseudomonas, Clostridium III, Acetivibrio*, members of the unclassified *Eubacteriaceae* and *Ruminococcaceae* and fungi loads were increased in infected S ponies, suggesting that strongyle and fungi may contribute to each other’s success in the ecological niche of the equine intestines. In contrast, butyrate-producing bacteria such as *Ruminococcus, Clostridium XIVa* and members of the *Lachnospiraceae* family decreased in S relative to R ponies. Additionally, these gut microbiota alterations induced changes in several immunological pathways in S ponies, including pathogen sensing, lipid metabolism, and activation of signal transduction that are critical for the regulation of immune system and energy homeostasis. These observations shed light on a putative implication of the gut microbiota in the intrinsic resistance to strongyle infection.

Overall, this longitudinal study provides a foundation to better understand the mechanisms that underpin the relationship between host susceptibility to strongyle infection, immune response and gut microbiota under natural conditions in horses and should contribute to the development of novel biomarkers of strongyle susceptibility and provide additional control options.

## 1. Introduction

Grazing horses are infected by a complex community of parasitic helminths, manly strongyles (Bucknell et al., 1995; Corning, 2009; Kuzmina et al., 2016; Ogbourne, 1976). Like other worms, strongyles have a direct life cycle in which they can survive outside of its host in pastures as well as in the horse’s intestines (Taylor et al., 2007). Infective larvae (L3 stage) are usually ingested and migrate to their preferred niche in the small or large intestine. After two molts, they will eventually become sexually mature adults and will lay eggs that are passed onto the pasture in the feces (Taylor et al., 2007).

Equine strongyle species are classified into *Strongylinae* and *Cyathostominae*, which differ, among other criteria, by their respective size (Lichtenfels et al., 2008). *Strongylus vulgaris* is the most pathogenic of the large strongyles as a result of the intestinal infarction that larval stages can cause during their migration. Its prevalence has been drastically reduced since the release of modern anthelmintics (Nielsen et al., 2012). On the one hand, nearly all horses are infected by small strongyles or cyathostomins throughout the world (Bucknell et al., 1995; Lyons et al., 1999; Ogbourne, 1976). Compared to *S. vulgaris*, small strongyles are responsible for milder symptoms, including weight loss or poor hair condition (Love et al., 1999). Infections are more common in immature animals meaning there is likely an immune component to infection susceptibility (Lyons et al., 1999). Furthermore, larval stages encyst into the colonic mucosa as part of their life cycle where millions can remain for years (Love et al., 1999; Matthews, 2014). Cyathostomin larvae encyst mostly in the autumn and winter in temperate zones of the northern hemisphere, accounting for up to 90% of the total worm burden (Matthews, 2014). The massive emergence of these larvae results in larval cyathostominosis, which is characterized by abdominal pain and diarrhea (Love et al., 1999). Treatment failure in this case can result in the death of horses in at least a third of cases (Giles et al., 1985), underscoring the need for efficient anthelmintic compounds.

Drug inefficacy reports have accumulated worldwide over the recent years and resistant isolates are now to be found in Europe (Geurden et al., 2014; Sallé et al., 2017), America (Smith et al., 2015) and Oceania (Scott et al., 2015). Additional control strategies are therefore required to alleviate selection pressure put on cyathostomin populations by anthelmintic treatments. One of the possible approaches is to exploit the over dispersion of strongyle infection in a herd to treat the only highly infected horses (Lester and Matthews, 2014). Indeed, it has been estimated that 80% of the total worm burden is produced by 20% of horses (Wood et al., 2012) and that 21% of the inter-individual variation had a heritable component (Kornaś et al., 2015). However, the factors underpinning this phenotypic contrast (Lester et al., 2013; Sallé et al., 2015) still remain unclear.

Equine strongyles are in close contact with a large community of microorganisms in the host intestines, estimated to reach a concentration of 10^9^ microorganisms per gram of ingesta in the cecum alone (Mackie and Wilkins, 1988), spanning 108 genera (Mach et al., 2017; Steelman et al., 2012; Venable et al., 2017) and at least seven phyla (Costa et al., 2012, 2015; Mach et al., 2017; Shepherd et al., 2012; Weese et al., 2015). Bacterial populations differ greatly throughout the various compartments of the equine gastrointestinal tract (*e.g.* duodenum, jejunum, ileum and colon) due to differences in the gut pH, available energy sources, epithelial architecture of each region, oxygen levels and physiological roles (Costa et al., 2015; Ericsson et al., 2016). The gut microbiota promotes digestion and nutrient absorption for host energy production and provides folate (Sugahara et al., 2015), vitamin K_2_ (Marley et al., 1986) and short chain fatty acids (SCFA) such as acetate, butyrate and propionate (Ericsson et al., 2016; Nedjadi et al., 2014). The gut microbiota also neutralizes drugs and carcinogens, modulates intestinal motility, protects the host from pathogens, and stimulates and matures the immune system and epithelial cells (reviewed by Nicholson et al. (2012)). Along with bacteria, both fungi and protozoa, comprise about 6-8% of the equine hindgut population (Dougal et al., 2013).

The physical presence of helminths in the intestinal lumen can alter the gut microbiota activity and composition (Midha et al., 2017; Peachey et al., 2017). These perturbations have been demonstrated in various host-nematode relationships including mice infected by *Heligmosomoides polygyrus* (Reynolds et al., 2014b; Su et al., 2017), *Nippostrongylus brasiliensis* (Fricke et al., 2015), or *Hymenolepis diminuta* (McKenney et al., 2015), ruminants (El-Ashram and Suo, 2017; Li et al., 2011), and pigs (Li et al., 2012; Wu et al., 2012), although no universal modification has been observed across systems. It also remains unresolved whether nematode infection has a beneficial (Lee et al., 2014) or a detrimental (Houlden et al., 2015) impact on the gut microbiota diversity, richness and functions. The mechanisms supporting these gut microbiota modifications also remain unclear and could arise indirectly because of the immune response helminths trigger in their host (Cattadori et al., 2016; Fricke et al., 2015; Reynolds et al., 2014b, 2015; Zaiss and Harris, 2016), such as regulatory T cell stimulation and lymphoid tissue modifications, alterations to the intestinal barrier (Boyett and Hsieh, 2014; Giacomin et al., 2016) or directly by the secretion of putative anti-bacterial compounds (Holm et al., 2015; Mcmurdie and Holmes, 2012) or modifications in the intestinal environment that supports them (D’Elia et al., 2009; Midha et al., 2017).

The putative interactions between equine strongyle infection and the gut microbiota and host physiology are unknown in horses. To address these questions, grazing ponies with extreme resistance or susceptibility toward natural strongyle infection were monitored over a five month period. We aimed to provide insights into the host response and the gut microbiota composition associated with strongyle natural infection that should guide the development of new microbiota-based control strategies.

## 2. Materials and methods

### 2.1. Animals selection

Twenty female Welsh ponies (10 resistant, R, and 10 susceptible, S; 5 ± 1.3 years old) from an experimental unit at the National Institute for Agricultural Research (INRA, UEPAO, Nouzilly, France) were selected from a set of 98 ponies monitored for fecal egg counts (FEC) since 2010. The FEC data were log-transformed to correct for over-dispersion and fitted a linear mixed model accounting for environmental fixed effects (month of sampling, year of sampling, time since last treatment, age at sampling). The individual was considered as a random variable to account for the intrinsic pony potential against strongyle infection. Estimated individual effects were centered and reduced to express each individual potential as a deviation from the mean on the logarithmic scale. Based on these values, two groups of the 10 most extreme ponies balanced for age were selected resulting in a resistant R group (mean individual effect with −1.18 phenotypic deviation from the mean and average age of 5.6 years) and a susceptible S group (mean individual effect with +1.45 phenotypic deviation from the mean and average age of 4.7 years). At inclusion, the median FEC was 800 eggs/g for the S group, whereas the median egg counts /g feces was 0 for the R ponies (Figure S1A, S1B and S1C).

As it has been previously established that gut microbiota profiles might be shaped by host genetics (Goodrich et al., 2014; Lozupone et al., 2012), this information was considered to understand the individual variance underlying microbiota composition. The kinship2 R package was used to create the genetic relationship matrix to estimate the genetic parameters and predicts breeding values between every considered pony. Both pedigree tree and correlation structure matrix are depicted in Figure S1D and Figure S1E, respectively.

All the procedures were conducted according to the guidelines for the care and use of experimental animals established by the French Ministry of Teaching and Research and the regional Val de Loire Ethics Committee (CEEA VdL, no 19). The protocol was registered under the number 2015021210238289_v4 in the experimental installations with the permit number: C371753. All the protocols were conducted in accordance with EEC regulation (n^o^ 2010/63/UE) governing the care and use of laboratory animals and effective in France since the 1^st^ of January 2013.

### 2.2. Longitudinal monitoring and sampling

The 20 Welsh ponies were treated with moxidectin and praziquantel (Equest Pramox®, Zoetis, Paris, France, 400 μg/kg of body weight of moxidectin and 2,5 mg/kg of praziquantel) in March 2015 to clear any patent and pre-patent infection and kept indoors for 3 months until the end of the moxidectin remanence period. They were maintained under natural light conditions in a 240 m^2^ pen with slatted floors, which precluded further nematode infections until they were moved to the experimental pasture. During housing, animals were fed with hay *ad libitum* and 600 g concentrate per animal per day. The concentrate (Tellus Thivat Nutrition Animale Propriétaire, Saint Germain de Salles, France) consisted of barley (150 g/kg), oat bran (162 g/kg), wheat straw (184.7 g/kg), oats (200 g/kg), alfalfa (121.7 g/kg), sugar beet pulp (50 g/kg), molasses (30 g/kg), salt (7.3 g/kg), carbonate Ca (5.5 g/kg) and a mineral and vitamin mix (2 g/kg), on an as-fed basis. The mineral and vitamin mix contained Ca (28.5%), P (1.6%), Na (5.6%), vitamin A (500,000 IU), vitamin D_3_ (125,000 IU), vitamin E (1,500 IU), cobalt carbonate (42 mg/kg), cupric sulfate (500 mg/kg), calcium iodate (10 mg/kg), iron sulfate (1,000 mg/kg), manganese sulfate (5,800 mg/kg), sodium selenite (16 mg/kg), and zinc sulfate (7,500 mg/kg) on an as-fed basis.

Grazing started by the end of a 3-month moxidectin remanence period (mid-June 2015). Ponies grazed from mid-June to the end of October 2015 at the Nouzilly experimental station (France). The experimental pasture (7.44 ha) consisted of tall fescue, *Festuca arundinacea;* timothy-grass, *Phleum prateonse*, meadow-grass, *Poa abbreviata;* soft-grass, *Holcus lanatus;* and cocksfoot, *Dactylis glomerata*). During all phases of the experimental period, ponies were provided *ad libitum* access to water.

Moxidectin is known to have moderate efficacy against encysted cyathostomin larvae (Reinemeyer et al., 2015; Xiao et al., 1994). Therefore, residual egg excretion can occur at the end of the remanence period, as a result of encysted larvae completing their development into adults. To eliminate this residual excretion and to avoid any interference with strongyle infection at pasture, a treatment targeting the only luminal immature and adult stages (pyrantel embonate; Strongid^®^ paste, Zoetis, Paris, France; single oral dose of 1.36 mg pyrantel base per Kg of body weight) was implemented at day 30.

Every pony was subjected to a longitudinal monitoring of fecal strongyle egg excretion and microbiota was performed monthly, *e.g.* 0, 24, 43, 92 and 132 days after the onset of grazing (Figure 1).

**Figure 1.**
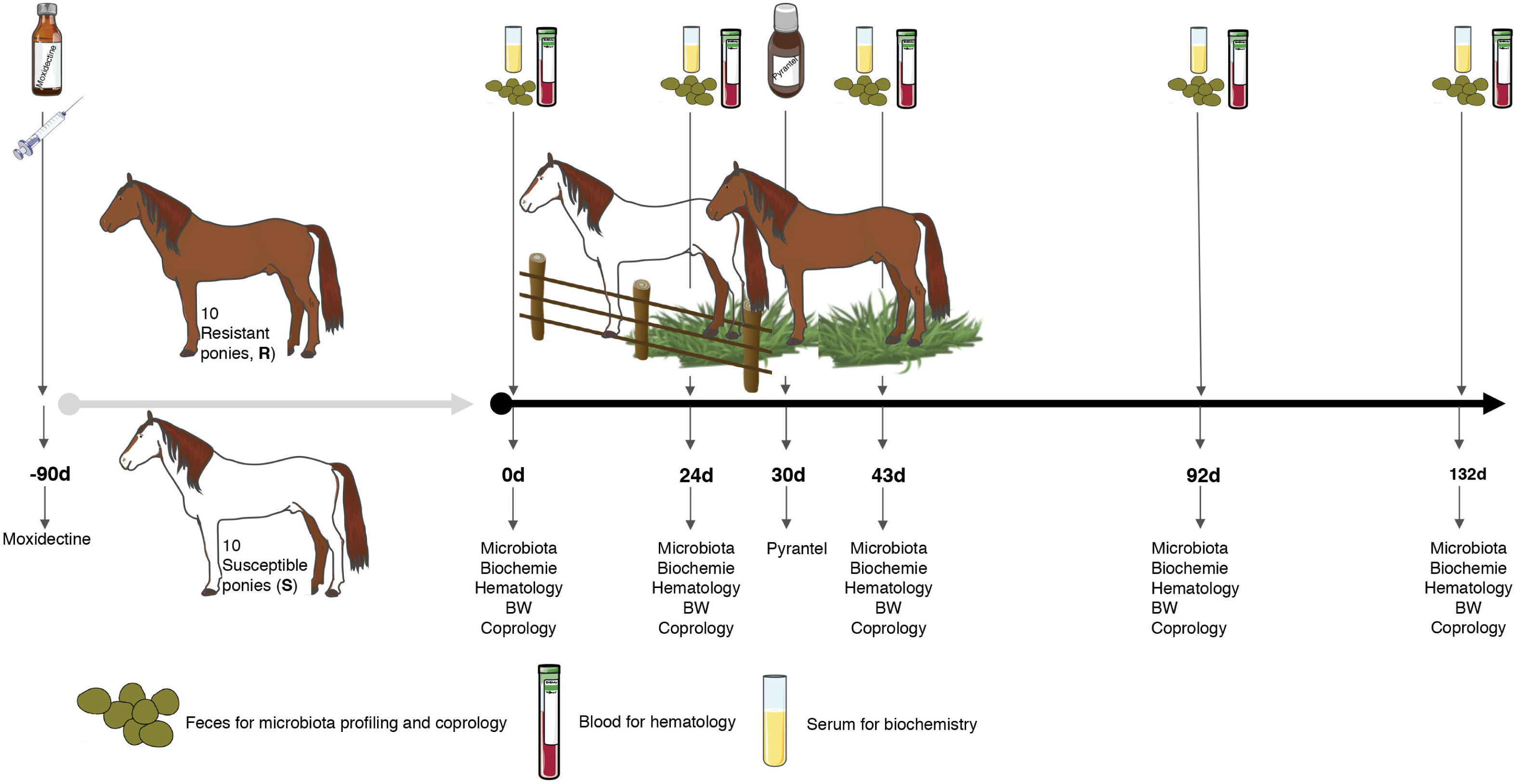
Experimental design and sampling. A set of twenty female ponies (10 susceptible (S) and 10 resistant (R) to strongylosis) were selected based on their fecal egg counts history during previous pasture seasons and were kept inside during the winter. In the spring, they were treated with moxidectin, to ensure that they were totally free from gastrointestinal nematodes (even from putative encysted larvae) and were kept indoors for three months. Thereafter, once no more effect of moxidectin treatment was detected, the ponies were moved to a 7.44 ha pasture to start the study. At day 30 of the study, a pyrantel treatment was administered to all the animals in order to reset the residual infections that could interfere in the protocol. A longitudinal monitoring of the parasitism level in each animal was performed through five time points from June to October. At each time point, fecal samples were collected from all ponies on 0, 24, 43, 92 and 132 days after the beginning of the grazing season to carry out fecal egg counts, pH measurements, and microbiota profiling. Blood samples were taken at the same time points to analyze biochemical and hematological parameters. This figure was produced using Servier Medical Art, available from www.servier.com/Powerpoint-image-bank

Fecal samples were collected from the rectum. Fecal aliquots for microbiota analysis were immediately snap-frozen in liquid nitrogen and stored at −80°C until DNA extraction, whereas fecal aliquots to measure the fecal egg counts were immediately sent to the laboratory. The pH in the feces was immediately determined after 10% fecal suspension (wt/vol) in saline solution (0.15 M NaCl solution).

Blood samples were taken from each pony and collected in EDTA-K3-coated tubes (5 mL) to determine hematological parameters and heparin tubes (10 mL) to determine biochemical parameters. After clotting, the heparin tubes were centrifuged at 4000 rpm during 15 min and the harvested plasma was stored at −20°C until analysis. Additionally, blood collected in EDTA-K3-tubes was used to measure the different blood cells.

For each individual, body weight and average daily weight gain were recorded until the end of the experiment. None of the ponies received antibiotic therapy during the sampling period and diarrhea was not detected in any ponies.

### 2.3. Fecal egg counts

Fecal egg counts (eggs per gram of wet feces) was measured as a proxy for patent strongyle infection. FEC was carried out using a modified McMaster technique (Raynaud, 1970) on 5 g of feces diluted in 70 mL of NaCl solution with a density of 1.2 (sensitivity of 50 eggs/g). A Wilcoxon rank-sum test with Benjamini-Hochberg multiple test correction was used to determine whether there was a significant difference between groups across the experiment. A *q* □< □0.05 was considered significant.

### 2.4. Blood hematological and biochemical assays

For blood hematological assays, blood were stirred at room temperature for good oxygenation during 15 min. Different blood cells were analyzed, including leucocytes (lymphocytes, monocytes, neutrophils, basophils and eosinophils), erythrocytes and different blood parameters such as hematocrit, mean corpuscular volume and the thrombocytes. The total blood cells were counted with a MS9-5 Hematology Counter^®^ (digital automatic hematology analyzer, Melet Schloesing Laboratories, France).

The serum biochemical parameters (albumin, cholesterol, globin, glucose, phosphatase alkaline, total proteins and urea) were measured colorimetric method using Select-6V rings with the M-ScanII Biochemical analyzer (Melet Schloesing Laboratories, France).

Mixed-effects analysis of the variance (ANOVA) or Wilcoxon rank-sum tests were conducted for continuous variables fitting a normal or non-normal distribution respectively to delineate whether there was a significant difference between the average values of phenotype traits for the different groups, using a significance level of *p* < 0.05. Blood cell counts were corrected for the mild dehydration by fitting the hematocrit as a co-variable in the model.

### 2.5. Weather data

Daily precipitation and temperatures were recorded at a meteorological station located 14 km from the experimental area.

### 2.6. Pasture contamination

Pasture contamination was assessed before the entry to the pasture and during the experiment. The number of infective larvae (L3) per kg of dry herbage was measured by 100 x 4 random sampling of grass on pasture following previously described method (Gruner and Raynaud, 1980). First, 600 g of grass were mixed with 20 mL of neutral pH soap diluted in 5L of tap water to allow larvae migration. This solution was subsequently left at room temperature prior to washing for larval recovery. Second, the washed material was passed through a coarse mesh sieve (20 cm of diameter) and collected in a container, then passed through both a 125 μm mesh sieve and a 20 μm mesh sieve. Third, the solution of the filtered material was split into four 10 mL glass tubes before centrifugation at 2500 rpm for 5 min. Water was soaked and replaced by a dense solution (NaCl, density 1.18-1.2). Lastly, a cover slip was added on the top of each. Tubes were subsequently centrifuged gently at 1500 rpm for 8 min to allow larval material adhere onto the cover slip, before further examination under an optical microscope. This step was performed four times.

### 2.7. Pasture chemical analysis

Hay was sampled at day 0 via grab samples from multiple depths into a bale and then composited. Herbage samples were collected from three randomly selected zones in the experimental pasture at days 24, 43, 92 and 132. At each sampling time, 10 hand-plucked samples simulating « bites » were taken from pasture. All samples were frozen until chemical analyses. Chemical compositions of samples were determined according to the “Association Française de Normalisation” (AFNOR) procedures: NF V18-109 (AFNOR, 1982) for dry matter (DM); NF V18-101 (AFNOR, 1977b) for crude ash; NF V18-100 (AFNOR, 1977a) for crude proteins; NF V03-040 (AFNOR, 1993) for crude fiber; NF V18-117 (AFNOR, 1977a) for crude fat; NF V18-122 (AFNOR, 1997) for neutral detergent fiber (NDF), acid detergent fiber (ADF) and acid detergent lignin (ADL). All of them were assayed with a heat stable amylase and expressed with exclusive of residual ash according to the method of Van Soest et al. (1991). Non-fiber carbohydrate (NFC), also called neutral detergent soluble carbohydrate (NDCS) were obtained by calculations: NFC = 1000 – CP – CF – Mm – NDF.

### 2.8. Microorganisms DNA extraction from feces samples

Total DNA was extracted from aliquots of frozen fecal samples (200 mg; 100 samples at different time points from 20 ponies), using E.Z.N.A.^®^ Stool DNA Kit (Omega Bio-Tek, Norcross, Georgia, USA). The DNA extraction protocol was carried out according to the manufacturer’s instructions (Omega-Bio-Tek, Norcross, Georgia, USA).

### 2.9. V3–V4 16S rRNA gene amplification

The V3-V4 hyper-variable regions of the 16S rDNA gene were amplified with two rounds of PCR using the forward primer (5’-CTTTCCCTACACGACGCTCTTCCGATCTACGGRAGGCAGCAG-3’) and the reverse primer (5’-GGAGTTCAGACGTGTGCTCTTCCGATCTTACCAGGGTATCTAATCCT-3’) modified in order to include Illumina adapters and barcode sequences which allow for directional sequencing. The first round of amplification was performed in triplicate in a total volume of 50 μL containing 10 ng of DNA, 2.5 units of a DNA-free Taq DNA Polymerase and 10X Taq DNA polymerase buffer (MTP Taq DNA Polymerase, Sigma). Subsequently, 10 mM of dNTP mixture (Euromedex, Souffelweyersheim, France), 20 mM of each primer (Sigma, Lezennes, France) and Nuclease-free water (Ambion, Thermo Fisher Scientific, Waltham, USA) were added. Ultrapure Taq DNA polymerase, ultrapure reagents, and plastic were selected in order to be DNA-free. The thermal cycle consisted of an initial denaturation step (1 min at 94°C), followed by 30 cycles of denaturation (1 min at 94°C), annealing (1 min at 65°C) and 1 min of extension at 72°C. The final extension step was performed for 10 min at 72°C. Amplicons were then purified using magnetic beads (Clean PCR system, CleanNA, Alphen an den Rijn, The Netherlands) as follows: beads/PCR reactional volume ratio of 0.8X and final elution volume of 32 μL using Elution Buffer EB (Qiagen). The concentrations of the purified amplicons were checked using a NanoDrop 8000 spectrophotometer (Thermo Fisher Scientific, Waltham, USA).

Sample multiplexing was performed thanks to 6 bp unique indexes, which were added during the second PCR step at the same time as the second part of the P5/P7 adapters used for the sequencing step on the Illumina MiSeq flow cells with the forward primer (5’-AATGATACGGCGACCACCGAGATCTACACTCTTTCCCTACACGAC-3’) and reverse primer (5’-CAAGCAGAAGACGGCATACGAGATNNNNNNGTGACTGGAGTTCAGACGTGT-3’). This second PCR step was performed using 10 ng of purified amplicons from the first PCR and adding 2.5 units of a DNA-free Taq DNA Polymerase and 10X MTP TaqDNA polymerase buffer (Sigma). The buffer was complemented with 10 mM of dNTP mixture (Euromedex), 20 mM of each primer (Eurogentec, HPLC grade) and Nuclease-free water (Ambion, Life Technologies) up to a final volume of 50 μL. The PCR reaction was carried out as follows: an initial denaturation step (94°C for 1 min), 12 cycles of amplification (94°C for 1 min, 65°C for 1 min and 72°C for 1 min) and a final extension step at 72°C for 10 min. Amplicons were purified as described for the first PCR round. The concentration of the purified amplicons was measured using Nanodrop 8000 spectrophotometer (Thermo Scientific) and the quality of a set of amplicons (12 samples per sequencing run) was checked using DNA 7500 chips onto a Bioanalyzer 2100 (Agilent Technologies, Santa Clara, CA, USA). All libraries were pooled at equimolar concentration in order to generate equivalent number of raw reads with each library. The final pool had a diluted concentration of 5 nM to 20 nM and was used for sequencing. Amplicon libraries were mixed with 15% PhiX control according to the Illumina’s protocol. Details on sequencing, PhiX control and FastQ files generation are specified elsewhere (Lluch et al., 2015). For this study, one sequencing run was performed using MiSeq 500 cycle reagent kit v2 (2x250 output; Illumina, USA).

### 2.10. Sequencing Data Preprocessing

Sequences were processed using the version 1.9.0 of the Quantitative Insights Into Microbial Ecology (QIIME) pipeline (Caporaso et al., 2010; Rideout et al., 2014) and by choosing the open-reference operational taxonomic units (OTU) calling approach (Rideout et al., 2014). First, forward and reverse paired-end sequence reads were collapsed into a single continuous sequence according to the ‘fastq-join’ option of the ‘join_paired_ends.py’ command in QIIME. The fastq-join function allowed a maximum difference within overlap region of 8%, a minimum overlap setting of 6 bp and a maximum overlap setting of 60 bp. The reads that did not overlap (~20% of the total) were removed from the analysis. The retained sequences were then quality filtered. De-multiplexing, primer removal and quality filtering processes were performed using the ‘split_libraries’_fastq.py command in QIIME. We applied a default base call Phred threshold of 20, allowing maximum three low-quality base calls before truncating a read, including only reads with >75% consecutive high-quality base calls, and excluding reads with ambiguous (N) base calls (Navas-Molina et al., 2013).

Subsequently, the sequences were clustered into OTUs against the GreenGenes database (release 2013-08: gg_13_8_otus) (DeSantis et al., 2006) by using the uclust (Edgar, 2010) method at a 97% similarity cutoff. The filtering of chimeric OTUs was performed by using Usearch version 6.1 (Edgar et al., 2011) against the GreenGenes reference alignment (DeSantis et al., 2006). A phylogenic tree was generated from the filtered alignment using FastTree (Price et al., 2010). Singletons were discarded from the dataset to minimize the effect of spurious, low abundance sequences using the ‘filter_otus_from_otu_table.py’ script in QIIME. To confirm the annotation, the resulting OTU representative sequences were then searched against the Ribosomal Database Project naïve Bayesian classifier (RDP 10 database, version 6 (Cole et al., 2009) database, using the online program SEQMATCH (http://rdp.cme.msu.edu/seqmatch/seqmatchintro.jsp). Finally, consensus taxonomy was provided for each OTU based on the taxonomic assignment of individual reads using GreenGenes and RDP databases. Using OTU abundance and the corresponding taxonomic classifications, feature abundance matrices were calculated at different taxonomic levels, representing OTUs and taxa abundance per sample. The “Phyloseq” (Mcmurdie and Holmes, 2012) and “Vegan” (Dixon, 2003) R package were used for the detailed downstream analysis on abundance matrix.

In the end, a total of 8,010,052 paired-end 250 bp reads were obtained, 6,428,315 of which were retained as high-quality sequences (Table S1). On average, a total of 58,015 sequences per sample were obtained in the study, with a mean length of 441 ± 15 bp. These sequences were clustered into 15,784 OTUs using the reference-based OTU-picking process (Table S2). Among them, 12,069 were classified taxonomically down to the genus level (Table S2). OTU counts per sample and OTU taxonomical assignments are available in Table S2. We filtered out unclassified taxa from the analysis because the main goal of the current study was to identify specific taxa related to host susceptibility to strongyle infection.

The α-diversity indexes (observed species richness, Chao1(Chao, 1984) and Shannon (Shannon, 1997)) were calculated using the “Phyloseq” R package (Mcmurdie and Holmes, 2012). Shannon’s diversity index is a composite measure of richness (number of OTUs present) and evenness (relative abundance of OTUs). The nonparametric Wilcoxon rank-sum test was used to compare α-diversity indexes between groups.

Relative abundance normalization was applied, which divides raw counts from a particular sample by the total number of reads in each sample.

To estimate β-diversity, un-weighted and weighted UniFrac distances were calculated from the OTU and genera abundance tables, and used in principal coordinates analysis (PCoA), correspondence analysis (CA), and non-parametric multidimensional scaling (NMDS) with the “Phyloseq” R package. The Permutational Multivariate Analysis of Variance (PERMANOVA), on un-weighted and weighted UniFrac distance matrices were applied through the Adonis function from “Vegan” R package to test for groups effect. In addition to multivariate analysis, we used the analysis of similarities (ANOSIM) to test for intragroup dispersion. ANOSIM is a permutation-based test where the null hypothesis states that within-group distances are not significantly smaller than between-group distances. The test statistic (*R*) can range from 1 to −1, with a value of 1 indicating that all samples within groups are more similar to each other than to any other samples from different groups. *R* is ≈0 when the null hypothesis is true, that distances within and between groups are the same on average.

The Wilcoxon rank-sum test with Benjamini-Hochberg multiple test correction was used to determine the differentially abundant OTUs, phyla, families, and genera between groups. A □ *q* □ < 0.25 was considered significant. This threshold was employed in previous microbiome studies because allows compensation for the large number of microbial taxa and multiple comparison adjustment (Lim et al., 2017).

This targeted locus study project has been deposited at DDBJ/EMBL/GenBank under the accession KBTQ01000000. The version described in this paper is the first version, KBTQ01000000. The bioproject described in this paper belongs to the BioProject PRJNA413884. The corresponding BioSamples accession numbers were SAMN07773451 to SAMN07773550.

### 2.11. Functional metagenomic predictions

The functional prediction for the 16S rRNA marker gene sequences was done using the phylogenetic investigation of communities by reconstruction of unobserved states (PICRUSt) (Langille et al., 2013). After excluding the unknown OTUs from the GreenGenes reference database and normalizing by 16S rRNA gene copy number, functional metagenomes for each sample were predicted from the Kyoto Encyclopedia of Genes and Genomes (KEGG) catalogue and collapsed to a specified KEGG level. We used Wilcoxon rank-sum test with Benjamini-Hochberg multiple test correction to evaluate pathway-level enrichments between groups. A □ *q* □ <0.05 was considered as significant.

### 2.12. Network inference at the genus level

Networks at the genus level were inferred between groups at different time points. In order to prevent the compositional effects bias typical of the classical correlations methods, we calculated the correlations among genera using the PCIT method, which identifies significant co-occurrence patterns through a data-driven methodology based on partial correlation and information theory as implemented in the PCIT algorithm (Reverter and Chan, 2008). Further details are depicted in Ramayo-Caldas et al. (2016). The genera with < 0.1% mean relative abundances were excluded to acquire the results for the taxa that met the statistical conditions for correlation estimations. Nodes in the network represent the genera and edges that connect these nodes represent correlations between genera. Based on correlation coefficient and *p-* values for correlation, we constructed co-occurrence networks. The cutoff of p-values was 0.05. The cutoff of correlation coefficients was determined as r≥|0.35|. Network properties were calculated with the NetworkAnalyzer plugin in Cytoscape. We used the “iGraph” R package to visualize the network. Strong and significant correlation between nodes (r≥|0.60|) were represented with larger edge width in the network.

### 2.13. Real-time quantitative PCR (qPCR) analysis of bacterial, fungal and protozoan loads

Loads of protozoa, anaerobic fungi and bacteria in fecal samples were quantified using a QuantStudio 12K Flex real-time instrument (Thermo Fisher Scientific, Waltham, USA). Primers for real-time amplification of ciliates, anaerobic fungi and bacteria have already been described in Mach et al. (2017) and have been purchased from Eurofins Genomics (Ebersberg, Germany).

Amplified fragments of the target genes were used and diluted 10-fold in series to produce seven standards, ranging from 2.25 × 10^7^ to 2.25 × 10^13^ copies per μg of DNA for bacteria and protozoa and ranging from 3.70 × 10^6^ to 3.70 × 10^12^ copies per μg of DNA for ciliates and fungi. Each reaction contained, in a final volume of 20 μL, 10 μL of Sybergreen Mix (Power SYBR Green PCR Master Mix, ThermoFisher, Ullkirch-Graffenstaden, France), 0.6 μM of each primer to final concentration of 300 mM, and 2 μL of standard or DNA template at 0.5 ng/μL. The primer concentration of anaerobic fungi was 200 mM and 150 mM for ciliate protozoa. The DNA template was 0.5 ng/μL. In all cases, the thermal protocol for qPCR amplification and detection included an initial step of denaturation of 10 min (95 °C), followed by 40 amplification cycles [15 s at 95 °C; 60 s at 60 °C]. After each run, melting curves between 60 and 95 °C were evaluated to confirm the absence of unspecific signals. For each sample and each gene, qPCR runs were performed in triplicate. The standard curve obtained the reference genomic fragment was used to calculate the number of copies of bacteria, protozoa or anaerobic fungi in feces. Taking into account the molecular mass of nucleotides and fragment length, we calculated the copy number as follows: mass in Daltons (g/mol) = (size of double-stranded [ds] product in base pairs [bp]) (330 Da × 2 nucleotides [nt]/bp). Wilcoxon rank-sum tests were calculated for all possible group combinations. A *p* < □0.05 was considered significant.

## 3. Results

The effects of natural strongyle infection on gut microbiota composition and host phenotypic variables were determined in ten resistant and ten susceptible grazing ponies over a five-month grazing season (Figure 1). Metagenomic, parasitological, hematological and biochemical measures were performed at five time points (Figure S2), hereafter referred to as days after the onset of the grazing season or grazing days (gd).

### 3.1. A mixture of abiotic and biotic environmental stressors during the 43-day transitioning period induced shifts in immunological and microbiota profiles

Measured FEC demonstrated a residual egg excretion in one susceptible individual at day 0 and in two susceptible ponies after 24 gd (Figure 2A and 2B). This was due to the known imperfect efficacy of moxidectin against encysted stages of strongyle. Because we were interested in studying the effects of strongyle exposure on the gut microbiota, a pyrantel treatment was administered after 30 gd to reset luminal parasite stages to zero in every pony. This treatment has a short-lived effect and resulted in negative FECs at 43 gd (Figure 2A and 2B). Therefore, the 0-43 gd period was considered as a transitioning period resulting in both, mild parasite exposure and changes in environmental conditions.

**Figure 2.**
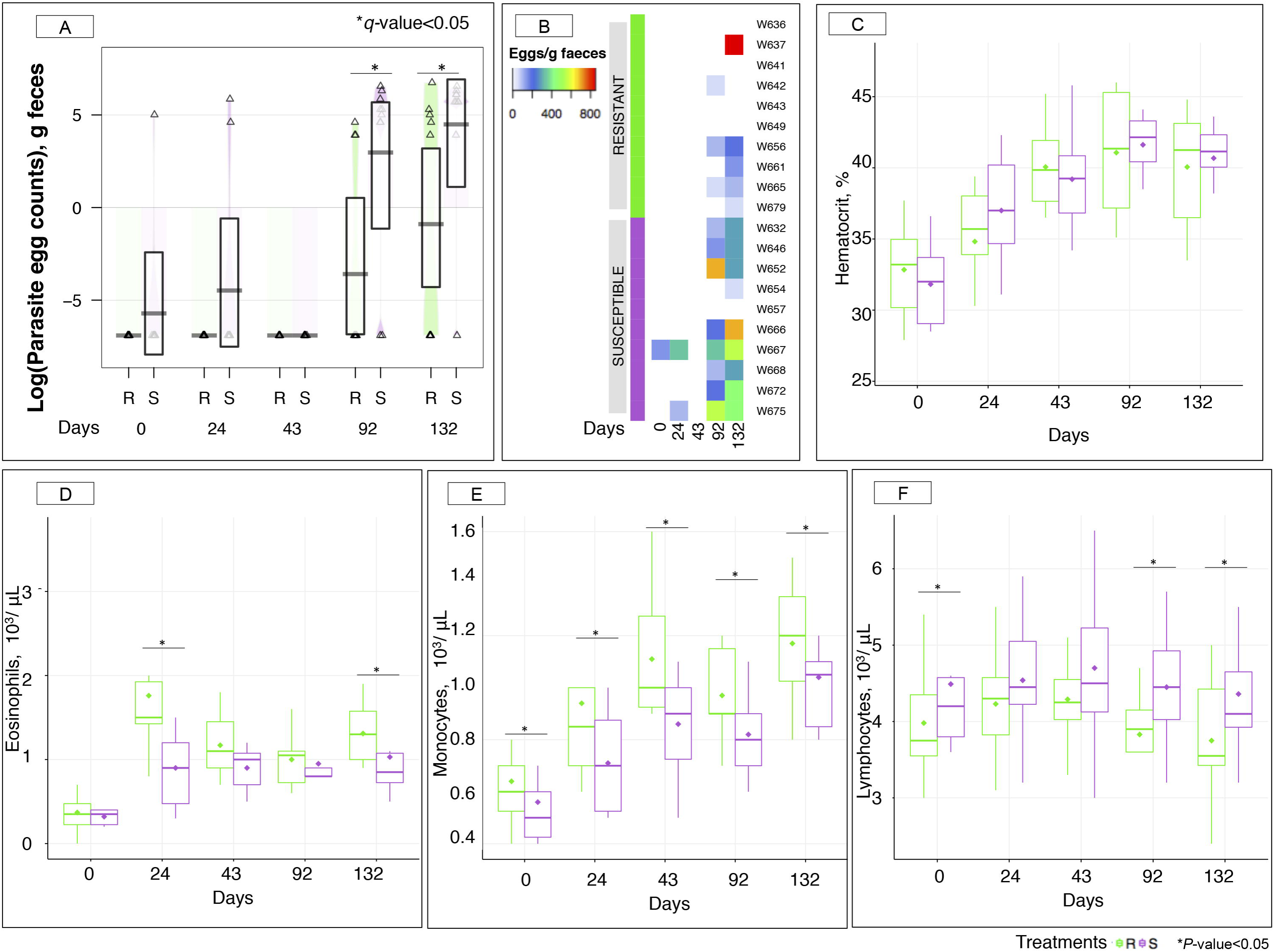
Fecal egg counts and hematological parameters between susceptible and resistant animals across time. (A) Boxplot of the log parasite fecal egg counts (eggs/g feces) in susceptible (S) and resistant (R) animals. Purple and green stand for S and R ponies respectively. *, *q* value < 0.05 for comparison between S and R ponies in each time point; (B) Heatmap of individual egg counts (eggs/g feces) in S and R animals (each row corresponding to one individual) across time (column). In the heatmap, egg count values range from 0 (white) and low (blue) to high values (red); (C) Hematocrit (%) between susceptible (S, violet boxes) and resistant (R, green boxes) animals across time. The quantification of different type of leukocytes: eosinophils (D), monocytes (E), and lymphocytes (F) were described between susceptible (S, violet boxes) and resistant (R, green boxes) animals across time. In all cases, boxes show median and interquartile range, and whiskers indicate 5th to 95th percentile. *, *p* value < 0.05 for comparison between S and R ponies in each time point.

Indeed, a heat wave took place over the first 43 days of the trial resulting in almost no rainfall (0.2 mm) and high temperatures peaking at 37.7 °C (Figure S3A). This situation ultimately led to grass senescence and lowered pasture quality (Figure S3B, Table S3). Mild dehydration also occurred in ponies as supported by a constant rise in measured hematocrits from 32.33 to 39.63 % during the first 43 gd (Figure 2C). To account for this, blood cell counts were corrected by hematocrit through time. Recorded hematological data showed that ponies were neither anemic nor thrombocytopenic throughout the experiment (Figure S4).

Concomitantly to these challenging conditions, increased levels of some white blood cell populations were observed, namely eosinophils (2.75-fold increase, *p* = 5.98 x 10^−13^, Figure 2D), and monocytes (1.64-fold increase, *p* = 2.94 × 10^−7^; Figure 2E) during the first 43 gd. Notably, R ponies had significantly higher levels of these immunological cells from 24 to 43 gd relative to S ponies. Grass samples analysis did not evidence any infective larvae until 43 gd.

16S rRNA gene sequencing was used to profile the fecal microbiota of the S and R ponies across time. Despite the distinctive susceptibility to parasite infection, the overall community structure showed no statistically significance difference in un-weighted (presence/absence) Unifrac analysis (PERMANOVA, p > 0.05) or abundance-weighted analysis (PERMANOVA, p > 0.05) during the first 43 days of the experiment. Measures of α-diversity (Chao1 richness, observed species and Shannon diversity index) were not significantly different between the two groups (Wilcoxon rank-sum test, *p* > 0.05). Although the overall gut microbial community of S ponies during the first 43 days of experiment was almost indistinguishable from those of R ponies, we next evaluated the association between the abundance of specific gut microbial taxa and the susceptibility to parasite infection. Statistical differences were not evident at phylum and family levels (Figure 3A). Only seven genera were statistically significant at a nominal *p* value < 0.05 (Wilcoxon rank-sum test; Figure 3B and 3C, Table S4) at day 0, including *Acetivibrio, Clostridium XIVa, Ruminococcus, Halella, Syntrophococcus*, unclassified *Lachnospiracea*, and *Lachnospiracea incertae sedis*, whereas a total of 20 taxa had differential presence at a nominal *p* value < 0.05 at day 24 (Wilcoxon rank-sum test; Figure 3C, Table S4). Specifically, species belonging to the *Blautia* and *Paraprevotella* genera were relatively more abundant (*p* < 0.05) in the S group compared to the R group (Table S4) at 24 gd. Conversely, other genera belonging to the Clostridiales order (*e.g. Clostridium sensu stricto, Clostridium IV, Clostridium III, Syntrophococcus, Oribacterium, Dehalobacterium, Mogibacterium, Acetivibrio, Sporobacter* and unclassified *Ruminococcaceae*) and Bacteroidales (*e.g. Hallella, Rikenella, Paraprevotella* and *Bacteroides*) were more abundant in the R group than in the S group at 24 gd (Figure 3B). These shifts in microbial taxa were not associated with modifications in functional gene abundances, as predicted from 16S rRNA data analysis (*q* > 0.05; Table S5).

**Figure 3.**
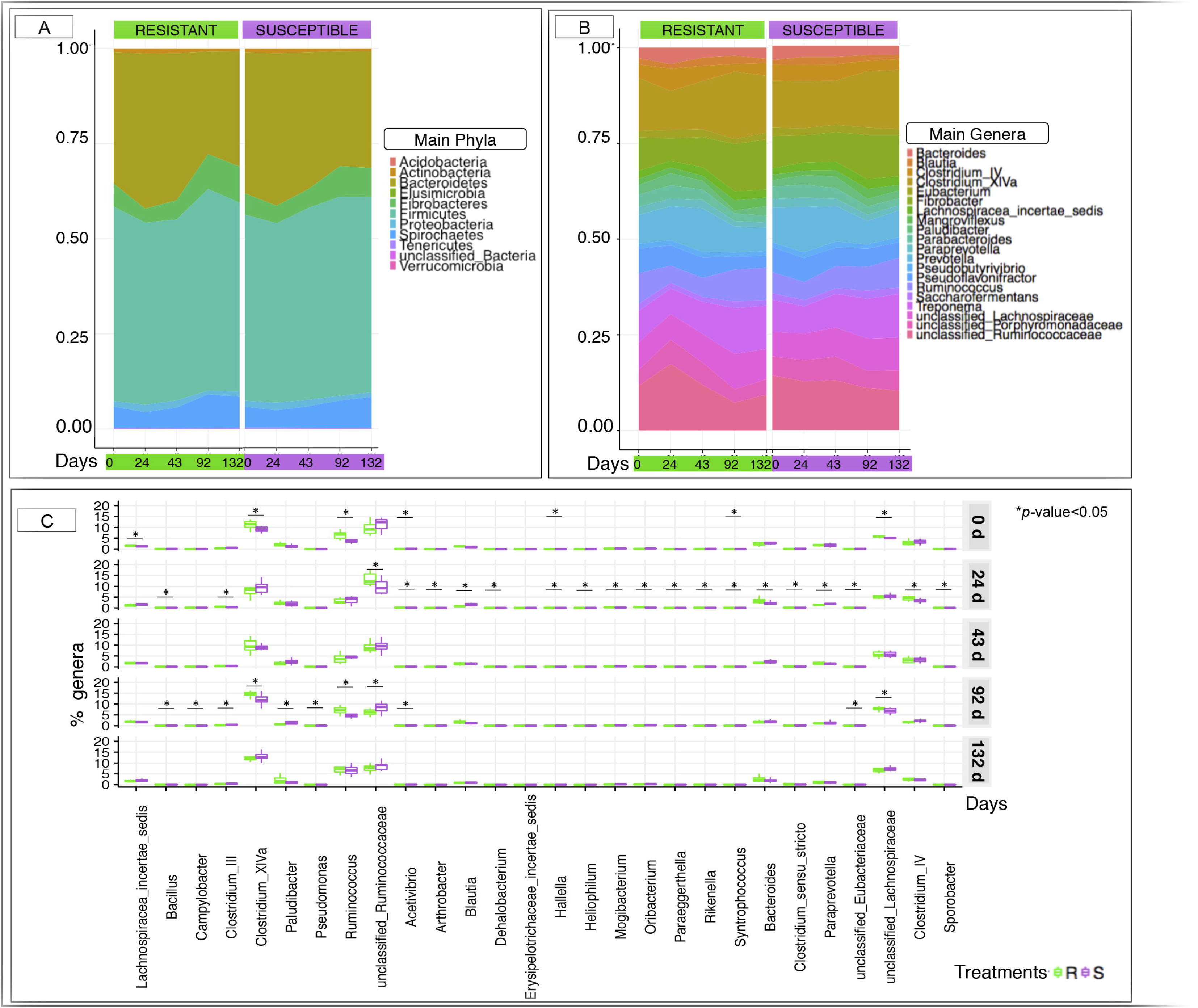
Dynamics of microbiota composition between susceptible and resistant animals across time. (A) Area plot representation of the phyla detected in feces between susceptible (S) and resistant (R) animals across time; (B) Area plot representation of the most abundant genera in feces between susceptible (S) and resistant (R) animals across time; (C) Boxplot graph representation of genera significantly affected between S and R animals across time. In all cases, susceptible animals are colored in violet and resistant animals in green. Boxes show median and interquartile range, and whiskers indicate 5th to 95th percentile. *, *p* value < 0.05 for comparison between S and R ponies in each time point.

### 3.2. The predicted levels of resistance matched observed fecal egg counts between resistant and susceptible ponies and shifts in the gut microbiota composition

By the end of the transition period, *e.g.* after 43gd, ponies were considered adapted to their new environmental conditions. The resetting of luminal stages with pyrantel resulted in negative FEC across ponies at 43 days after the onset of grazing (Figure 2A and 2B). After the patent infection, parasite egg excretion was significantly higher in the susceptible group after 92 (235 eggs in S and 20 eggs in R ponies on average) and 132 gd (340 eggs in S and 135 eggs in R ponies on average; Figure 2A).

The ponies’ body weights (Figure S5A) and average daily weight gains (Figure S5B) did not show significant differences between groups after natural parasite infection, and none of them displayed clinical symptoms like lethargy or diarrhea. However, strongyle exposure induced contrasted shifts in white blood cell populations between the two groups of ponies. Circulating monocyte levels were higher (*p* < 0.05) in R in comparison to S ponies through the whole period (Figure 2E). But the opposite trend was found for circulating lymphocytes, which were significantly enriched in the white blood cells population in S ponies during parasite infection (Figure 2F). Among the white blood cell population, R ponies also presented higher levels of eosinophils at 132 gd (*p* < 0.05, figure 2D). Similarly, serum biochemical analyses revealed a mild elevation in albumin, cholesterol, the enzyme alkaline phosphatase and total proteins from 43 gd to the end of the experiment in both groups, as well as elevated levels of urea, in particular at 92 and 132 gd (Figure S6).

Strongyle mediated alterations in microbiota diversity and structure were investigated between the two groups of ponies. As for the transition period, both groups of ponies displayed equivalent microbial species richness and alpha-diversity indexes (Wilcoxon rank-sum test, *p* > 0.05, Figure 4A). The diversity Chao1 and Shannon indexes were similar between the two groups through time (Wilcoxon rank-sum test, *p* > 0.05, Figure 4B). The UniFrac distance followed by PCoA (Figure 4C) showed no distinct clustering between samples from the S and the R group, which was indicative of, if at all, minor differences in microbiota composition between the two groups of ponies during parasite infection. Similarly, the correspondence analysis (Figure 4D) and the Jaccard network (Figure 4E) analyses suggested that the overall gut microbiota composition was largely similar between S and R ponies at each time point.

**Figure 4.**
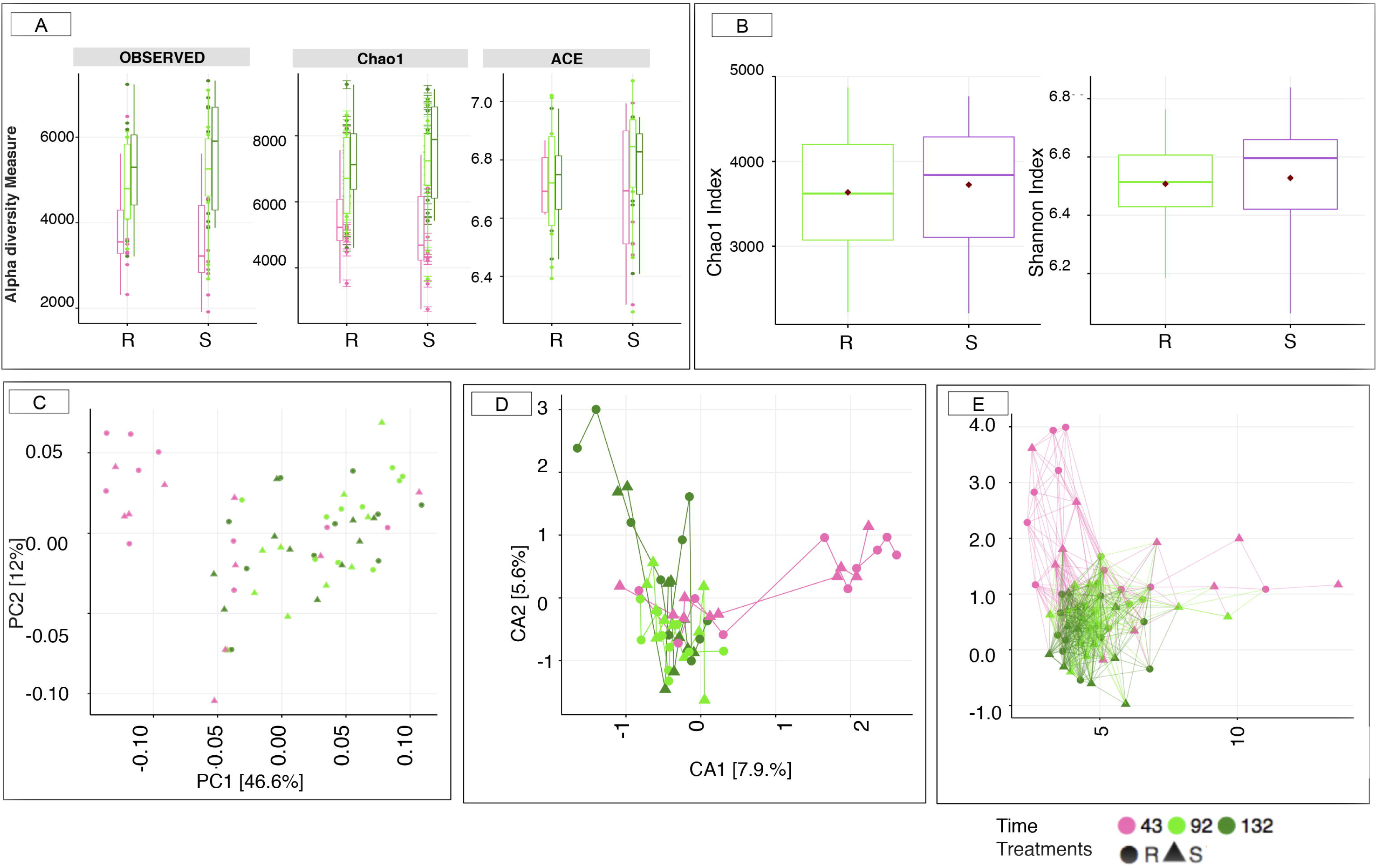
Estimation of the α-diversity indexes and ß-diversity insusceptible and resistant animals during the natural parasite infection (from day 43 to day 132). (A) Estimation of the α-diversity indexes in susceptible (S) and resistant (R) during the natural parasite infection (from day 43 to 132). The box color indicates the time point analyzed: (pink=43 d, green=92 d, and darkgreen=132 d); (B) α-diversity indexes between S and R animals during the natural parasite infection. Susceptible animals were colored in violet and resistant animals in green; (C) Principal Coordinate analysis of Unifrac distances to compare fecal communities at the level of genera that differ between S and R animals across natural parasite infection. Both PC axes 1 and 2 were plotted. Together they explained 58.6% of whole variation; (D) Correspondence analyses of Unifrac distances to compare fecal communities at the level of genera that differ between S and R animals across natural parasite infection. Both CA axes 1 and 2 were plotted; (E) Genus-level network representation between ponies across natural parasite infection linked within a specified Jaccard distance of 0.85. Two samples were considered “connected” if the distance between them was less than 0.85. In all cases, the relative position of points was optimized for the visual display of network properties. The point’s shape indicates the susceptibility to strongylosis (triangle: susceptible (S); round: resistant (R)), the node color indicates the time point analyzed: (pink=43 d, green=92 d, and darkgreen=132 d).

However, changes in relative abundance of certain genera concomitantly arose with strongyle egg excretion at 92 gd. For example, *Paludibacter*, *Campylobacter, Bacillus, Pseudomonas, Clostridium III, Acetivibrio*, and members of the unclassified family *Eubacteriaceae* and *Ruminococcaceae* increased in S relative to R ponies (*p* < 0.05 and *q* ≤ 0.25; Figure 3C). Moreover, the relative abundance of *Acetivibrio*, and *Clostridium* III highly correlated with FEC (Pearson correlation coefficient ρ > 0.60). This genera enrichment was concomitant with depletion in *Ruminococcus, Clostridium XIVa* and members of the *Lachnospiraceae* family (*p* < 0.05 and *q ≤* 0.25, Figure 3C). The complete list of differentially expressed genera and q-values is presented in Supplementary Table S4. These modifications in the gut bacterial community structure between S and R also resulted in functional modifications, as inferred from PICRUSt. Noteworthy, among the well-characterized bacterial functions, S ponies microbiota tended to show an enrichment (*q* < 0.10) of the mineral absorption, protein digestion and absorption, as well as of some of the pathways related to cell motility (*e.g.* bacterial chemotaxis, bacterial motility proteins, flagellar assembly), lipid metabolism (sphingolipid metabolism), peroxisome, and signal transduction (phosphatidylinositol signaling system) among others compared to the R group at 92 gd (Table S5).

Because the magnitude and direction of changes for *Clostridium XIVa, Ruminococcus, Acetivibrio* and unclassified *Lachnospiracea* observed at 92 gd between the two groups were already remarked at day 0, we evaluated the host genetics. Interestingly, 80% of the ponies from group R presented higher genetic relatedness (Figure S1E), whereas there was no evidence for high genetic relatedness between S individuals.

The gut microbiota co-occurrence networks were marginally different between S and R ponies at 92 gd (Figure S7). The topological properties were calculated to describe the complex pattern of inter-relationships among nodes, and to distinguish differences in taxa correlations between these two groups of ponies (Table S6). The structural properties of the S network were slightly greater than the R network, indicating more connections and closer relationships of microbial taxa in the S group. Notably, S co-occurrence network displayed higher levels of betweenness centrality, which measures the number of shortest paths going through a given node, and higher degree levels, which describes the number of neighbors relative to R network. Nodes with the highest degree and betweenness centrality values were identified as key genera in the co-occurrence networks. The keystone genera in S were related to the Clostridiales order (*e.g. Clostridium IV, Roseburia, Nakamurella*) or Spirochaetales (*e.g. Treponema*), whereas R network keystone included members of Clostridiales order (*e.g. Clostridium XIV, Roseburia*) and Bacteroidales (*e.g Alloprevotella*).

To understand the modifications of the gut environment after the natural infection, feces pH measurements and the fungal, protozoan and bacterial loads in feces were investigated between groups of ponies. There were no differences in pH between the groups at any time point, although feces pH significantly decreased after the onset of patent strongyle infection (Figure S8A). Interestingly, anaerobic fungal loads were higher in the S than in the R group (*p* < 0.05) with more than 0.5 log of difference at day 43 (Figure S8B), while protozoan loads were lower in S group than in the R group at the same time point (Figure S8C). Bacteria loads were constant throughout the experiment (Figure S8D).

### 3.3. Gut microbiota composition and functions shifted immediately after natural parasite infection

Data of both groups were pooled together to assess the influence of natural strongyle infection (from day 43 to day 132) on gut microbiota composition, irrespective of the ponies’ predicted susceptibility. Analysis of similarities tests demonstrated that microbiota at day 43 were highly dissimilar and significantly divergent from day 92 and day 132 (R=0.197, *p* < 0.001). Interestingly, species richness increased significantly at day 92 and remained high until the end of the experiment (Figure 5A). Similarly, PCoA, CA and Jaccard network demonstrated a high variability in the distribution of microbiota between day 43 and the other points across the natural parasite infection (Figure 4). Consequently, the most significant alterations at genus level were found at 92 gd relative to 43 gd, which included a decrease (*q* < 0.05; Figure 5B) in the relative abundances of members of the order Bacteroidales (*e.g. Alloprevotella, Petrimonas, Paludibacter*), Clostridiales (*e.g. Clostridium IV, Oscillibacter, Ethanoligenens*) and Proteobacteria (*e.g. Desulfovibrio*). On the other hand, the relative abundances of dominant genera such as *Clostridium XIVa, Fibrobacter, Ruminococcus, Treponema* and of the members of the as yet unclassified family *Lachnospiraceae* were significantly increased (*q* < 0.05, Figure 6B). The complete list of increased and decreased genera including direction, coefficient and *q*-values is presented in Supplementary Table S7. These temporal changes on gut microbiota between day 43 and day 132 correlated with an increase in strongyle egg counts and an improvement in pasture quality with less non-digestible carbohydrate components and more N and mineral content (Figure S2A).

**Figure 5.**
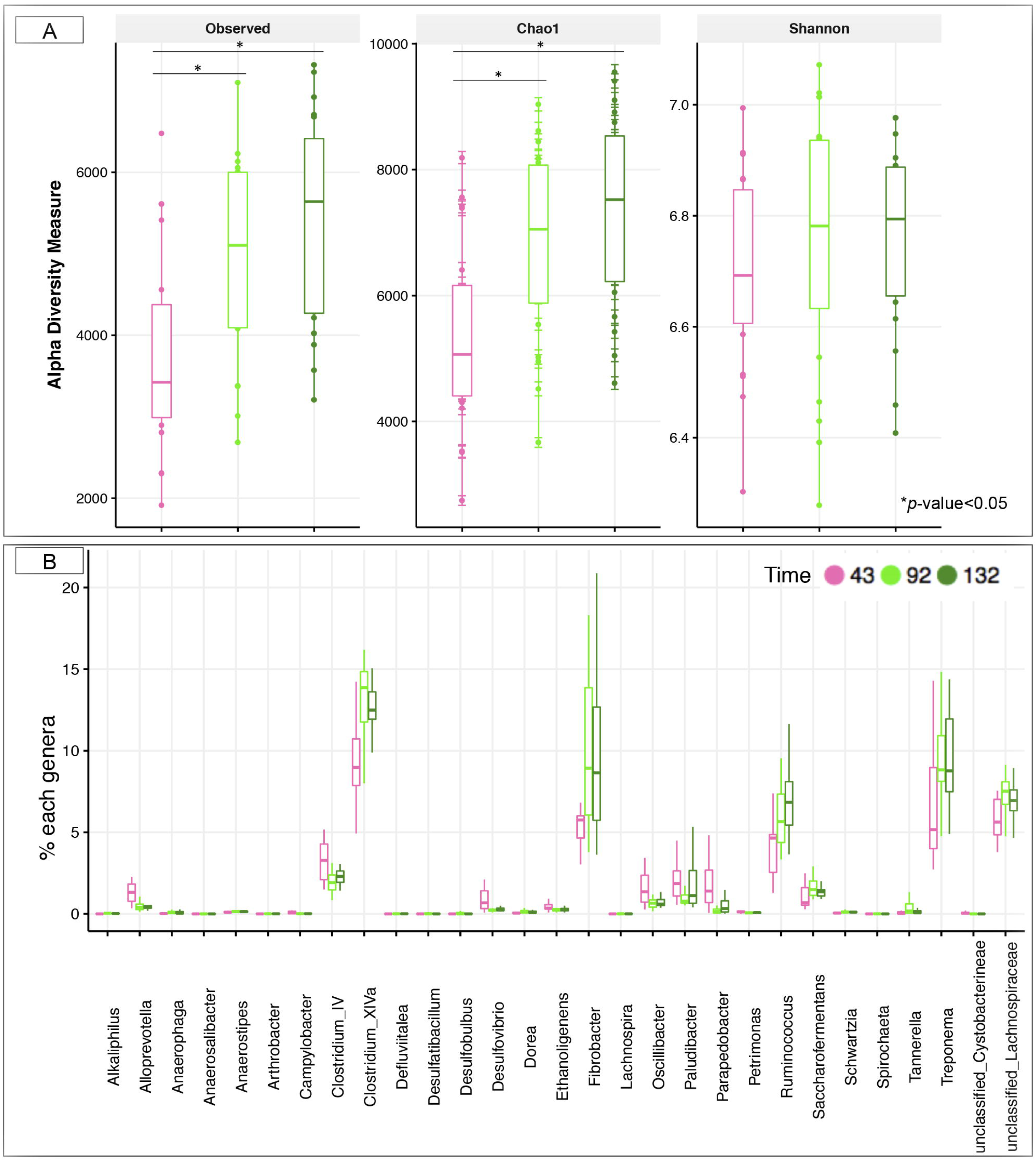
Longitudinal dynamics of microbiota composition upon natural parasite infection. (A) Estimation of the α-diversity indexes across the natural parasite infection: from day 43 to day 132. The box color indicates the time point analyzed: (pink=43 d, green=92 d, and darkgreen=132 d); (B) Boxplot graph representation of genera significantly affected (*q* < 0.05) at day 92 relative to day 43. In all cases, the box color indicates the time point analyzed: pink=43 d, green=92 d, and darkgreen=132 d. Boxes show median and interquartile range, and whiskers indicate 5th to 95th percentile. All genera plotted were statistically significant (*q* < 0.05) between day 92 and day 43.

These microbiota alterations had an effect on a broad range of biological functions. A total of 94 pathways displayed significantly different abundance (50% related to metabolism) between 43 and 92 gd. Interestingly, bacterial invasion of epithelial cells was among the top enriched pathways at day 92 (*q* < 0.0003; Table S8). Conversely, only 34 pathways were found to be different between 92 and 132 gd. Similarly, 50% of the different metabolic potentials were related to metabolism, including amino acid, energy, carbohydrate, lipid and xenobiotic metabolism.

## 4. Discussion

While alternative control strategies are needed for a more sustainable control of horse strongyle infection, the factors contributing to the over-dispersed distribution of these parasites in their hosts remain poorly characterized (Debeffe et al., 2016; Kornaś et al., 2015; Wood et al., 2012). In their preferred niche, strongyles are surrounded by gut microbiota and reciprocal interactions between them are expected. Our study aimed to identify the consequences of parasite infection on the gut microbiota and host physiology under natural conditions and to seek for a metagenomics signature of strongyle infection in resistant and susceptible ponies. Notably, R and S ponies showed contrasted immune responses toward natural strongyle infection, and their gut microbiota displayed variations at the genus level. Additionally, we showed that besides the host susceptibility to strongyle infection, variations in gut microbiota occurred after the onset of strongyle egg excretion during, expanding earlier observations on the association between parasites and gut microbiota composition on other species (Aivelo and Norberg, 2017; Cooper et al., 2013; Ramanan et al, 2016; El-Ashram and Suo, 2017; Fricke et al., 2015; Houlden et al., 2015; Li et al., 2011, 2012; McKenney et al., 2015; Newbold et al., 2017; Osborne et al., 2015; Reynolds et al., 2014b; Su et al., 2017; Walk et al., 2010; Wu et al., 2012; Zaiss et al., 2015; Zaiss and Harris, 2016). Moreover, for the first time we reported the effect of intestinal parasites have on the gut microbiota in horses.

Under our experimental setting, the combined heat wave and the associated reduced pasture yield and quality induced a mixture of different stresses during the first 43 days of the experiment. Under these challenging climatic and nutritional conditions, R ponies displayed higher levels of eosinophils and monocytes from 24 to 43 gd in comparison to the S ponies. Eosinophilia has been reported in experimentally challenged horses (Murphy and Love, 1997). However, no larvae were recovered from pasture samples in our study suggesting mild, if any, contamination, in line with the drought conditions that are detrimental to their survival (Nielsen et al., 2007). In addition, the mild residual egg excretion observed after 24 grazing days occurred in only two susceptible ponies, which cannot explain the increased eosinophils in the R group. Therefore, this differential profile in immune cell populations may result from the stressful environmental conditions as already reported elsewhere (Collier et al., 2008). Remarkably, the contrast between R and S ponies was also found while comparing their respective gut microbiota composition. Although the constituent phyla and genera within the gut microbiota of R and S ponies were congruent with other studies based on horses (Costa et al., 2012, 2015; Mach et al., 2017; Shepherd et al., 2012; Steelman et al., 2012; Venable et al., 2017; Weese et al., 2015), *e.g.* members of Firmicutes, Bacteroidetes, Spirochaetes and Fibrobacteres predominating, R ponies presented an increase of several Clostridiales and Bacteroides species at day 24, whereas only species related to *Blautia* and *Paraprevotella* genera were relatively more abundant in the S. Indisputably, individuals with different susceptibility to parasite infection adapt to environmental stress in different ways. Whether the gut microbiota differences are ascribed to divergence in the immune response or are due to impaired nutrient availability remains unclear, but micronutrient deficiencies might dictate microbial-microbial as well as microbial-environmental interactions through the gut (Mach and Clark, 2017).

The most interesting findings brought forward by this work were obtained during the natural strongyle infection, from day 43 to 132 of the experiment (Figure 6A). Congruent with our initial hypothesis, the observed FEC matched the predicted resistance levels throughout the trial, hence supporting the high reproducibility of FEC (Debeffe et al., 2016; Scheuerle et al., 2016) and the feasibility to select for more resistant individuals (Kornaś et al., 2015). Observed FEC were associated with differing immune responses characterized by higher eosinophil and monocyte counts in R ponies and increased levels of circulating lymphocytes in S ponies during the infection, suggesting a direct functional relationship between parasite infection and immune response (Howitt et al., 2016). The induction of lymphocytes in S animals could have been a way to neutralize invading L3 and facilitate repair and turnover of injury tissue. In R, the eosinophilia, which is a well-recognized immune response to strongyle infection in horses (Reynolds et al., 2012) and plays an important role in destroying parasites by acting as a killer cell against larvae (Herbert et al., 2000), likely played a role in parasitic infection resistance (Lyons et al., 2000). However, the interpretation of the relationship between strongyle infection and immune profile after natural parasite infection in our study required accounting for likely confounding effects of yearly variation in environmental conditions (*e.g.* extreme heat temperatures and reduction of pasture quality).

**Figure 6.**
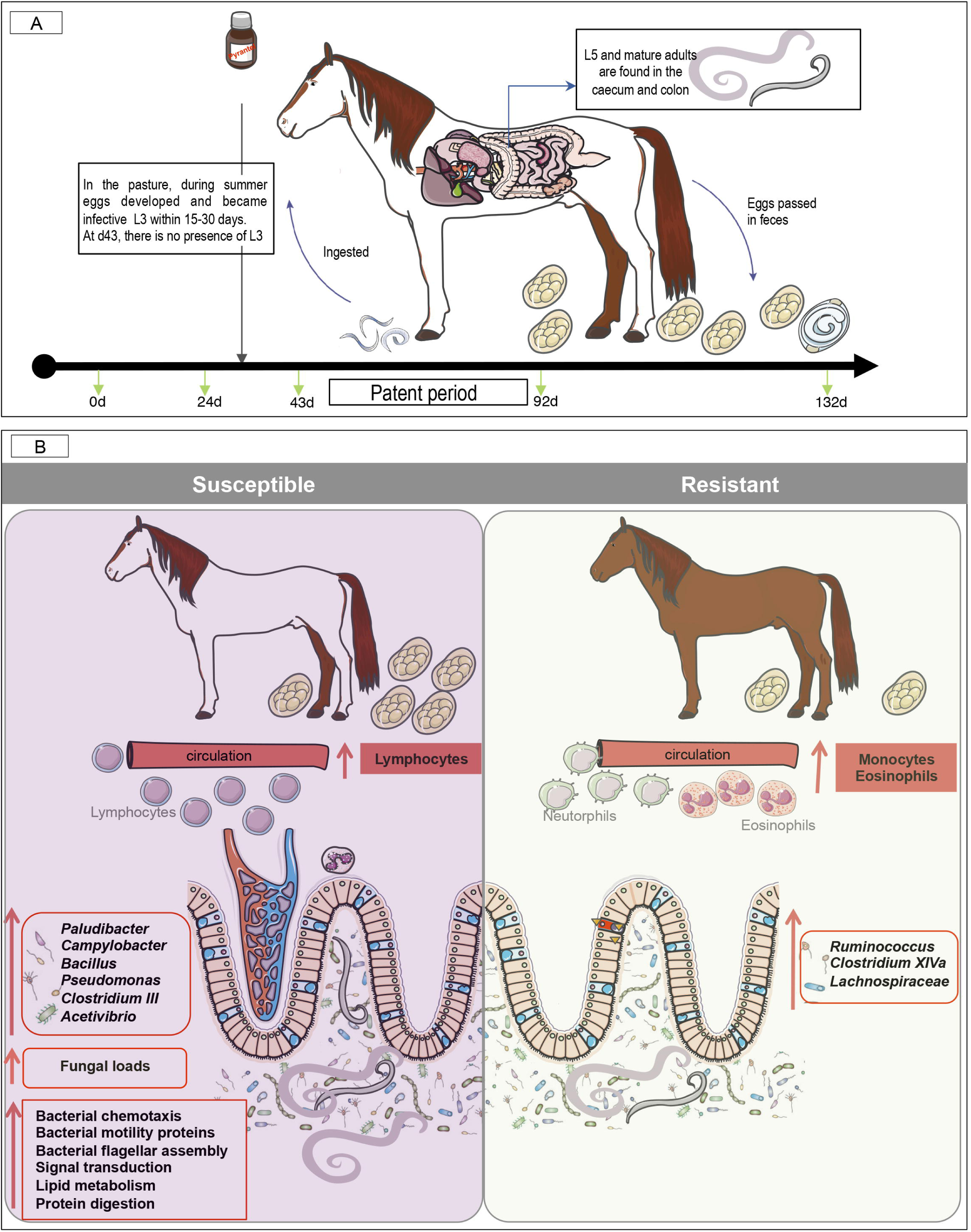
A model for gut microbiota modifications and their effects on host physiology after natural strongyle infection. We hypothesize that natural parasite infection in susceptible ponies increases the release of lymphocytes but decreases the monocytes and eosinophils cell counts. Concomitantly, parasite infection induced alterations in bacterial-fungal inter-kingdom, increasing the abundance of *Paludibacter, Campylobacter, Bacillus, Pseudomonas, Clostridium III, Acetivibrio* and the overall loads of fungi and parasite egg counts in the feces. On the other hand, butyrate producing bacteria such as members of *Ruminococcus, Clostridium XIVa* and *Lachnospiraceae* family were found to be depleted in susceptible ponies, but enriched in resistant animals, suggesting a possible effect of N-butyrate on the protection of inflammation in resistant animals. Because butyrate is a potent inhibitor of inflammation, it is suggested that susceptible ponies are prone to the gut inflammation because of the altered abundance of butyrate-producing bacteria. The lower N-butyrate bacteria abundance was accompanied by a number of microbiota functional pathways that reflected immunological mechanisms, including pathogen sensing, changes in lipids, and activation of signal transduction pathways inside of the cell. This figure was produced using Servier Medical Art, available from www.servier.com/Powerpoint-image-bank

Despite the limited infection level monitored throughout the trial, FEC differences between S and R ponies from day 92 to the end of the experiment were also reflected in the composition and function of their gut microbiota. Strongyle natural infection in S ponies coincided with an increase in pathobionts, such as *Pseudomonas, Campylobacter,* and *Bacillus,* anaerobic fungi loads as well as a reduction of commensal genera such as *Clostridium XIVa, Ruminococcus,* and unclassified *Lachnospiraceae* (Figure 6B). Firmicutes belonging to the families *Ruminococcaceae* (also referred as clostridial cluster IV) and *Lachnospiraceae* (also referred as clostridial cluster XIVa) comprise most of the butyrate-producing bacteria in the human gut (Geirnaert et al., 2017). Due to butyrate’s anti-inflammatory properties, it might be suggested that the higher helminth infection in susceptible ponies alters the abundance of butyrate-producing bacteria which therefore modulates the gut inflammation (Li et al., 2016). Additionally, the reduced abundance of *Clostridium XIV* in S ponies could have had functional importance for immune gardening against the overgrowth of pathobionts such as *Pseudomonas* and *Campylobacter. Clostridium XIVa* is one of the main methane-producing bacteria that has been implicated in the maintenance of mucosal homeostasis and protection against intestinal inflammatory diseases through the promotion of T_reg_ cell accumulation (Atarashi et al., 2011). In addition, it was notable that a significant number of microbiota functional pathways in S ponies reflected immunological mechanisms, including pathogen sensing, changes in lipids, and activation of signal transduction pathways inside of the cell that are critical for regulation of immune system and maintaining energy homeostasis (Vassart and Costagliola, 2011).

Altogether, our data suggest that parasitic infections in S ponies increased the risk of subsequent pathobionts overgrowth in the gut by reducing butyrate producing bacteria that may play an important role in mediating interactions between the host immune system and intestinal parasites (Oliver et al., 2003). Although the mechanisms of the interactions between these specific gut genera and the host require further elucidation, our findings suggest that specific modulations of the gut microbiota might be an effective strategy for managing parasite infections in horse.

The concomitant increase in fungi abundance in S ponies during the natural parasite infection, could have had an important pathophysiological consequence, especially in host fiber metabolism (Dougal et al., 2013) and inflammation (Noverr and Huffnagle, 2004). Fungi have been found to play a significant role in degrading cellulose and other plant fibers making them more accessible to bacteria (Güiris et al., 2010; Leng et al., 2011). This results in a consequent increase in SCFA such as acetate, butyrate and propionate (Ericsson et al., 2016; Nedjadi et al., 2014) which contribute to 60-70% of energy for horses (Al Jassim and Andrews, 2009). Consequently, the fungi-mediated increase in SCFA of microbial origin could have compensated for the decreased abundance of *Ruminococcus* in S ponies. The key contribution of *Treponema,* a non-pathogenic carbohydrate metabolizer (Han et al., 2011), to the co-occurrence network of S ponies at day 92 also suggests that compensatory mechanisms were induced to degrade fiber and supply the host with micro and macronutrients. More detailed nutritional determinations are needed to resolve this issue. Fungi can also secrete inflammatory substances that have been shown to reduce Treg cell responses and may elicit a Th17 innate immune responses and pro-inflammatory cytokines secretion (reviewed by Underhill and Iliev (2014)). In line with the seemingly symbiotic relationship between *H. polygyrus* and bacteria from the family *Lactobacillaceae* in mice (Reynolds et al., 2015), it could be speculated that strongyle and fungi may contribute to each other’s success in the ecological niche of the equine intestines.

Interestingly, R and S ponies exhibited some contrasting phenotypes from the beginning of the experiment under worm-free conditions. Indeed, the predicted R ponies displayed higher levels of circulating monocytes and lower lymphocytes than S ponies at day 0, but they also exhibited higher *Clostridium XIVa, Ruminococcus* and unclassified *Lachnospiracea* abundances and lower abundance of *Acetivibrio.* Because 80% of the ponies from group R presented higher genetic relatedness, we could not exclude the possibility that these specific taxa were influenced by host genetic factors and could sign the intrinsic resistance potential of ponies to strongyle infection. As explained above, *Clostridium XIVa* has been implicated in the maintenance of mucosal homeostasis and the protection against intestinal inflammatory diseases through the promotion of Treg cell accumulation (Atarashi et al., 2011). Therefore, it is tempting to speculate that R and S ponies had an intrinsic different modulation of the mucosal and systemic immunity as well as the gut bacteria composition and function before any infection took place.

Beyond the effects of susceptibility or resistance to strongyle natural infection on gut microbiota composition, our findings are consistent with other studies showing that helminths have the capacity to induce or maintain higher gut microbiota diversity (Giacomin et al., 2016; Lee et al., 2014; Newbold et al., 2017). In addition, we observed that the patterns of gut microbial alterations during parasite infection were overall highly consistent with other studies (Li et al., 2016; McKenney et al., 2015; Reynolds et al., 2014b; Su et al., 2017; Walk et al., 2010). Specifically, the expansion of Clostridiales has been already reported (Ramanan et al., 2016; McKenney et al., 2015; Walk et al., 2010; Zaiss et al., 2015) and the reduction of *Oscillibacter* have been found in pigs infected by *Trichuris suis* (Li et al., 2012; Wu et al., 2012). Nevertheless, we caution that the late microbiota changes from day 43 to day 92 could also reflect the effects of pasture quality improvements. Therefore, we hypothesize that the increased shifts in gut microbiota composition during parasite infection were partially explained by the nematodes inducing an anti-inflammatory environment and diverting immune responses away from themselves (Cattadori et al., 2016; Fricke et al., 2015; Reynolds et al., 2014b, 2015; Zaiss and Harris, 2016). This could also be explained by the changes in the microbial-microbial interactions, the contribution of increased levels of N availability and the non-fibrous carbohydrates in the pasture, as well as changes in the microbial-environmental interactions throughout the gut (D’Elia et al., 2009; Midha et al., 2017). In fact, concomitantly to the parasite infection, pH and fungi loads decreased and protozoa loads increased. As a result, survival and proliferation of certain microbial species become favored or depleted.

In conclusion, we showed that host parasite susceptibility correlated with parasite burden, with susceptible ponies having higher egg excretions than resistant animals throughout the experiment. However, due to the low level of helminth infections observed under natural conditions, differences in the immune response and gut microbiota composition between susceptible and resistant animals were modest. Eosinophils and monocytes populations were more abundant in resistant ponies while lymphocytes were less abundant in their blood, which may provide a health benefit relative to susceptible animals. Moreover, susceptible ponies presented a reduction of butyrate-producing bacteria such as *Clostridium XIVa, Ruminococcus,* and unclassified *Lachnospiraceae,* which may induce a disruption of the maintenance of mucosal homeostasis, intestinal inflammation and dysbiosis. In line with this hypothesis, an increase in pathobionts such as *Pseudomonas, Campylobacter* and fungi loads were observed in susceptible ponies. Our results therefore suggest that susceptibility to strongyle infection occurs in the presence of host genetic and other innate and gut environmental factors that influence immune response and affect individual risk. This investigation should be followed by experimental work in order to establish the causative reasons for variation in the microbiota.

## Authors and Contributors

AB, GS and NM designed the experiment. AC, GS and NM drafted the main manuscript text. NM designed and carried out the bioinformatics and biostatistical analyses, prepared all the figures and provided critical feedback on content. VB performed the RT-qPCR analyses. FR was in charge of pony maintenance and care throughout the experiment and managed sampling. AM analyzed the chemical composition of the diet. JC and CK performed fecal egg counts. MR performed the blood analysis. All authors reviewed the manuscript and approved the final version.

## Acknowledgements

We are grateful to Jean Marie Yvon, Yvan Gaude, Thierry Blard, Thierry Gascogne, Philippe Barriere, Francois Stieau and Adelaide Touchard for the animal sampling and management, and to Marine Guego, Marine Beinat, and Julie Rivière for participating to the sample collection during the project. We also thank Diane Esquerré for preparing the libraries and performed the MiSeq sequencing. Lastly, we are grateful to the INRA MIGALE bioinformatics platform (http://migale.jouy.inra.fr) for providing computational resources.

## List of abbreviations

ADF: acid detergent fiber
ADL: acid detergent lignin
AFNOR: Association Française de Normalisation
ANOSIM: analysis of similarities
ANOVA: Analysis of the variance
CA: correspondence analysis
DM: dry matter
FEC: fecal egg counts
KEGG: Kyoto Encyclopedia of Genes and Genomes
NDCS: neutral detergent soluble carbohydrate
NDF: neutral detergent fiber
NFC: Non-fiber carbohydrate
NMDS: non-parametric multidimensional scaling
OTU: operational taxonomic units
1PCoA: principal coordinates analysis
PERMANOVA: Permutational Multivariate Analysis of Variance
PICRUSt: phylogenetic investigation of communities by reconstruction of unobserved states
QIIME: Quantitative Insights Into Microbial Ecology
qPCR: real time quantitative PCR
RDP: Ribosomal Database Project naïve Bayesian classifier
SCFA: short chain fatty acids

## Supplementary Material

### Supporting tables

**Table S1. Summary of study samples and fecal bacterial 16S rRNA gene amplicon sequence datasets**

**Table S2. The OTU taxonomical assignments and OTU counts in each pony across time Table S3. Chemical composition of pasture across time.**

**Table S4. Differences in relative abundance of genera between susceptible (S) and resistant (R) animals across time.**

**Table S5. Differences in KEGG pathway abundance between susceptible (S) and resistant (R) animals across time.** Pathways values across time are represented as %, where normalized counts from a particular pathway is divides by the total number of counts in each sample.

**Table S6. Topological properties of co-occurring networks obtained from susceptible and resistant animals at day 92 of the experiment.**

**Table S7. Differences in relative abundance of genera from day 43 to day 132, regardless of susceptibility to strongyles.**

**Table S8. Differences in KEGG pathway abundance from day 43 to day 132, regardless of susceptibility to strongyles.** Pathways values across time are represented as %, where normalized counts from a particular pathway is divides by the total number of counts in each sample.

### Supporting figures

**Figure S1. Genetic kinship between the 20 Welsh ponies in the experiment**

(A) Heatmap of the historical egg counts (eggs/g feces) in susceptible (S) and resistant (R) animals included in the study. The fecal egg counts values of 4 continuous years were included. In the heatmap, rows represent the animals and columns the different time points analyzed. Not all animals have information for each time point. In the heatmap, red = high values, green = low values of egg counts per g of feces; (B) Violin plot of the historical fecal egg counts of the S and R animals included in the study. The median egg counts /g feces were 800 for S (the mean was 897, the first quartile was 350, and the third quartile was 1,375), whereas the median egg counts /g feces were 0 for R (the mean was 246.1864, the first quartile was 0, and the third quartile was 150); (C); Density plot of historical fecal egg counts values in S and R animals; (D) Pedigree plot. A six-generation pedigree plot is illustrated, with different shapes for male (squares) and female (circles). The shapes are black for the 20 ponies in the study; (E) Heatmap of the kinship coefficient matrix, which assess the genetic resemblance between ponies. Each entry in the matrix is the kinship coefficient between two subjects. Animals are arranged in the order of their genetic relatedness; genetically similar animals are near each other. Note that the diagonal elements did not have values above unity, showing no consanguineous mating in the families. The treatment (susceptible (S) or resistant (R) is delineated below to the animal name. In the heatmap, red = high values of genetic relatedness, white = low values of genetic relatedness.

**Figure S2. Overview of the data analysis in the study**

(A) Effect of parasite susceptibility on gut microbiota composition, parasite egg excretion, gut-related parameters and host parameters across time.

Step 1: Measurement of the gut microbiota composition between susceptible (S) and resistant (R) ponies across time. This step involved the analysis of the α-diversity and ß-diversity between S and R ponies across time, as well as the analysis to assess gut genera whose relative abundances changed between groups across time, the determination of the corresponding KEGG pathways and the inference of the co-occurrence network.

Step 2: Measurement of the gut related parameters between S and R ponies across time. The gut parameters included the pH, the fungal, bacteria and protozoan loads, as well as the number of parasite egg in the feces.

Step 3: Measurement of the host parameters between S and R ponies across time. The host parameters included the body weight, the daily average gain, hematological and biochemical parameters in blood.

**Figure S3. Whether and pasture quality throughout the experiment**

(A) Daily precipitation and temperatures recorded at a meteorological station located 14 km from the experimental pasture. Maximum and minimum temperatures are colored in red and blue, respectively. Dashed line represents precipitation; (B) Chemical composition (crude protein (CP), neutral and acid detergent fiber (NDF and ADF), crude fiber (CF), crude ash, and acid detergent lignin (ADL)) of hay at day 0 and pasture at day 24, 43, 92 and 132. From day 1 to day 43, the lack of rainfall resulted in significant soil moisture deficits and reduced growth rates of pasture. This period coincided with the late-flowering stage of the pasture, when stems and leaves are being depleted of nutrients and herbage maturation and lignification increases. After day 43, the environmental conditions were eminently favorable to start a second pasture cycle, with a quick herbage growth, high protein content and lower fiber content. We observed that herbage protein peaked at day 92, after the intense fall rains and the increased senescence of green material.

**Figure S4. Hemogram data in susceptible and resistant animals across time.**

The evaluation of the hemogram involved the determination of the hematocrit, total white blood cell counts, total erythrocyte count, erythrocyte indices and platelet counts. The determination of the hematocrit (A) and the total white blood cells (B) was performed between susceptible (S, violet boxes) and resistant (R, green boxes) animals across time. The quantification of different type of leukocytes: eosinophils (C), monocytes (D), lymphocytes (E), neutrophils (F) and basophils (G) were described between S and R animals across time. The values corresponding to red blood cell distribution width (H), microcytic (I) and macrocytic (J) platelets, as well as microcytic red blood cells (K), and macrocytic red blood cells (L) were plotted. The red blood cell distribution width is plotted in (L). In all cases, boxes show median and interquartile range, and whiskers indicate 5th to 95th percentile. *, *p* value < 0.05 for comparison between S and R ponies in each time point.

**Figure S5. Performance between susceptible and resistant animals across time**

(A) Boxplot and violin plot representation of body weight (kg) between susceptible (S) and resistant (R) ponies across time; (B) Boxplot and violin plot representation of average daily gain (kg) between S and R ponies across time. In all cases, susceptible animals were colored in violet and resistant animals in green.

**Figure S6**. **Biochemical data in susceptible and resistant animals across time.**

Levels of albumin (A), cholesterol (B), globins (C), glucose (D), alkaline phosphatase (E), ratio albumin/goblins (F), total proteins (G) and urea (H) in susceptible (S, violet boxes) and resistant animals (R green boxes) were plotted. Boxes show median and interquartile range, and whiskers indicate 5th to 95th percentile. *, *p* value < 0.05 for comparison between S and R ponies in each time point.

**Figure S7. Co-occurrence network at 92 days after the entry to the pasture for susceptible and resistant ponies**

The correlations among genera were calculated using the PCIT method, which identifies significant co-occurrence patterns. The size of the node is proportional to genera abundance. Node fill color corresponds to phylum taxonomic classification. Edges color represent positive (red) and negative (blue) connections, the edge thickness is equivalent to the correlation values. Only genera with a relative abundance > 0.10 were included.

**Figure S8. Microbiota functional parameters between susceptible and resistant ponies across time.**

(A) Boxplot graph representation of pH in feces between susceptible (S) and resistant (R) animals at different time points; (B) Boxplot graph representation of loads of anaerobic fungi in feces between S and R animals at different time points; (C) Boxplot graph representation of loads of protozoan in feces between S and R animals at different time points; (D) Boxplot graph representation of loads of bacteria in feces between S and R animals at different time points. In all cases, susceptible animals are colored in violet and resistant animals in green. *, *p* value < 0.05 for comparison between S and R ponies in each time point.

